# SenseNet, a tool for analysis of protein structure networks obtained from molecular dynamics simulations

**DOI:** 10.1101/2021.06.28.450194

**Authors:** Markus Schneider, Iris Antes

**Author notes:** Corresponding author (IA). **Software Availability**: All analyses were performed with our Cytoscape 3 plugin SenseNet, which is freely available at https://cytoscape.org/. The data preparation script AIFgen, the SenseNet manual and tutorials are available at https://bioinformatics.wzw.tum.de/.

## Abstract

Computational methods play a key role for investigating allosteric mechanisms in proteins, with the potential of generating valuable insights for innovative drug design. Here we present the SenseNet (“Structure ENSEmble NETworks”) framework for analysis of protein structure networks, which differs from established network models by focusing on interaction timelines obtained by molecular dynamics simulations. This approach is evaluated by predicting allosteric residues reported by NMR experiments in the PDZ2 domain of hPTP1e, a reference system for which previous computational predictions have shown considerable variance. We applied two models based on the mutual information between interaction timelines to estimate the conformational influence of each residue on its local environment. In terms of accuracy our prediction model is comparable to the top performing model published for this system, but by contrast benefits from its independence from NMR structures. Our results are complementary to experimental data and the consensus of previous predictions, demonstrating the potential of our new analysis tool SenseNet. Biochemical interpretation of our model suggests that allosteric residues in the PDZ2 domain form two distinct clusters of contiguous sidechain surfaces. SenseNet is provided as a plugin for the network analysis software Cytoscape, allowing for ease of future application and contributing to a system of compatible tools bridging the fields of system and structural biology.

**Author Summary:** Regulation and signal transduction processes in proteins are often correlated to structural changes induced by ligand binding, which can lead to suppression or enhancement of protein function. A common method to investigate such changes are numerical simulations of protein dynamics. We developed the analysis software SenseNet for predicting how protein dynamics and function is affected by e.g. ligand binding events based on molecular dynamics simulations. Our model estimates which structural elements of the protein confer the most information about their local environment, reasoning that these elements are essential for signal propagation. Applying this method on the PDZ2 domain of the hPTP1e protein, we were able to accurately predict structure elements with known signaling roles as determined by previous experiments. Integrating these experimental data with the consensus of other computational models and our predictions, we find two separate pathways which may transmit information through the PDZ2 protein structure. In addition to deepening our insight into the behavior of this particular protein, these results demonstrate the usefulness of our methods for other systems, such as potential drug targets. To make this analysis available to a broad audience, we implemented it as a plugin for the popular network analysis software Cytoscape.

## Introduction

Protein structure networks map atoms from a protein structure to nodes and define edges to represent atom interactions, e.g. contacts and hydrogen bonds. The resulting networks may be used to predict e.g. allosteric communication pathways (1-3) with potential applications in innovative drug design (4-8). Most commonly, such analyses are based on individual crystal structures and rely on centrality measures such as betweenness centrality (BC) or characteristic path length centrality (CPLC) to identify functionally important residues (1-3, 9). However, application of these algorithms to experimental structures of e.g. the PDZ domain did not provide results consistent with experiment (10). It has been generally recognized that highly dynamic effects such as allostery, which are not always associated with stable conformations, are difficult to study solely on the basis of individual experimentally obtained structures (5, 11-13). Computational methods for analyzing structure ensembles obtained from e.g. molecular dynamics simulations (MD), which capture the dynamic behavior of proteins, are therefore attractive for allosteric prediction (11, 14-20). Several tools exist for analysis of structure ensemble networks, among them xPyder (21), PyInteraph (22), MD-TASK (23), gRINN (24), PSN-Ensemble (25), NAPS (26, 27), RIP-MD (28), Bio3D (29), MDN (30) and the Cytoscape plugin RINalyzer (31). A common approach for network analysis of MD data is to define edges by correlation analysis of atomistic motions, which comes at the cost of losing structural and conformational details of the underlying interactions. In addition, many approaches use a rigid mapping of one node per residue, preventing the combination of different levels of resolution, e.g. to separate information flow between backbone and sidechain atoms. Finally, the majority of tools are provided as standalone programs or webservers, making it difficult to combine different algorithms within a single analysis session. To address these limitations, we developed SenseNet, a plugin for the free network analysis software Cytoscape (32). SenseNet is based on an alternative strategy to scalar correlation coefficients, namely associating edges with MD-based timelines, which allow to track the evolution of interactions during a simulation by checking their existence at predefined timeslots. This representation allows for a larger variety of analyses than correlation-based approaches, like e.g. interaction averages, lifetime analysis, frame clustering, or shared information between timelines.

Ligand binding often modulates protein function by triggering conformational changes distant from the binding site. A major goal of computational allosteric prediction is to identify key residues sensing ligand binding events over long intramolecular distances; in the context of computational predictions, these residues are commonly labeled as “allosteric”. For the purpose of evaluating these methods, PDZ domains are a well-established reference system. Members of this abundant domain class commonly bind C-terminal or short internal peptide sequences and participate in allosteric interactions with other domains (33, 34), serving as initiators and mediators of protein assembly processes (35-37). Although the domain is allosterically modulated by its peptide ligands, crystal and solution NMR structures of the PDZ2 domain of hPTP1e (human Protein-Tyrosine Phosphatase 1e) show no substantial conformational changes between apo and ligand bound states (38). Therefore, the relationship between structure, dynamics, and allostery in the PDZ2 domain of hPTP1e was explored by Lee and coworkers, who identified a number of allosteric residues by probing the effects of ligand binding and point mutations on NMR backbone and methyl side chain dynamics (38-40). However, open questions remain concerning the contribution of residues lacking methyl groups and how individual residues act together to form allosteric pathways, motivating structure-based computational prediction as a complementary strategy (41). Methods previously applied to the PDZ2 system include interaction energy and correlation networks (42, 43), elastic network models (44), hydrogen bond heat diffusion pathways (45), relative entropy networks of distance distributions (REDAN) (46), and coordinate fluctuations (47, 48). Furthermore, specialized simulation techniques were employed such as perturbation response scanning (49), rigid residue scan (RRS) (50), and NMR guided simulations (10, 51). However, results reported by computational studies have shown considerable variance, warranting efforts to consolidate and improve prediction models (41).

In this work, we present our network analysis software SenseNet and evaluate two of therein implemented, timeline-focused algorithms to find pathways of allosteric information transfer in the PDZ2 domain. By quantifying how much information the timelines of physical interactions provide about their environment, we obtained accurate models for predicting allosteric residues in PDZ2. Finally, we propose a consolidated allosteric model combining our results with experimental data and the consensus of previous predictions, which suggests that PDZ2 contains two allosteric pathways formed by clusters of contiguous sidechain surfaces.

## Materials & Methods

### Algorithms

In a structure network as implemented in SenseNet, each node (which together form the set of nodes *N*) represents a single atom or a group of atoms while edges represent interactions between nodes. If several interaction types (e.g. contacts or hydrogen bonds) are present, a node pair may be connected by more than one edge. First, we define an atomistic timeline as the vector

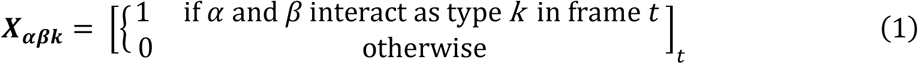

where *α, β* are nodes representing single atoms, *k* is an interaction type and *t* is a simulation time frame (bold type face denotes matrices and vectors). Timelines of edges connecting two atom groups (e. g. residues) are calculated as

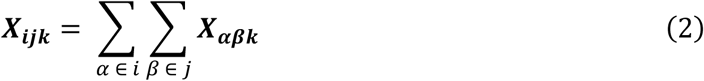

in which *i, j* are nodes representing atom groups. The connectivity between nodes is given by the symmetric adjacency matrix

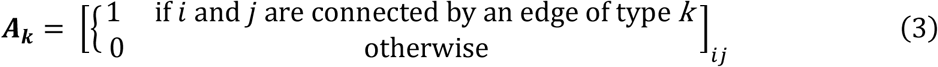

for each interaction type *k*. To describe the correlation between interactions, we define the edge neighbor correlation factor (ECF) as

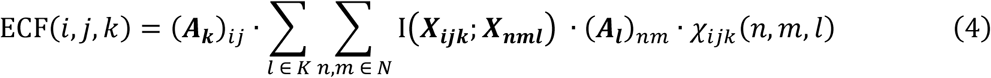

with *i, j* belonging to the node set *N, k* and *l* being part of the interaction type set *K*, and *I* is the mutual information function

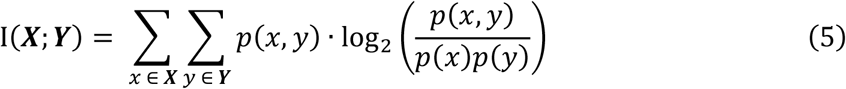

in which *p*(*x, y*) represents the joint probability of values *x* and *y* and *p*(*x*) corresponds to the marginal probability of value *x* in timeline *X*. Furthermore, *χ* represents an indicator function selecting the neighboring edges of *i, j, k* and is defined as

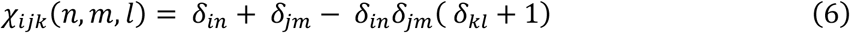

where *δ* is the Kronecker delta and the *δ*_*kl*_ term serves to exclude the self-information of edge *i, j, k* (Fig 1). Summing up the ECF scores of a node’s adjacent edges gives the node correlation factor

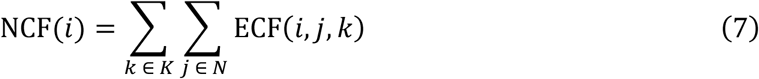

highlighting residues with strong conformational correlation with their environment. To quantify correlation changes between two simulations, after selecting one as analysis target and the other as reference, the definition of eq. 5 may be adjusted to

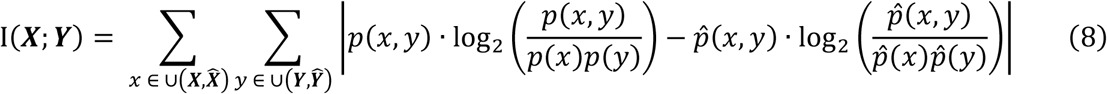

with 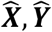 denoting the timelines from the reference simulation corresponding to *X* and *y* of the target simulation and 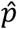 representing the probabilities of the reference timelines. Substitution of eq. 8 in eq. 4 yields the difference node correlation factor (DNCF). Note that edges which exist solely in the reference network do not contribute to the DNCF, therefore the score is not symmetric with respect to interchanging target and reference networks.

**Fig 1.**
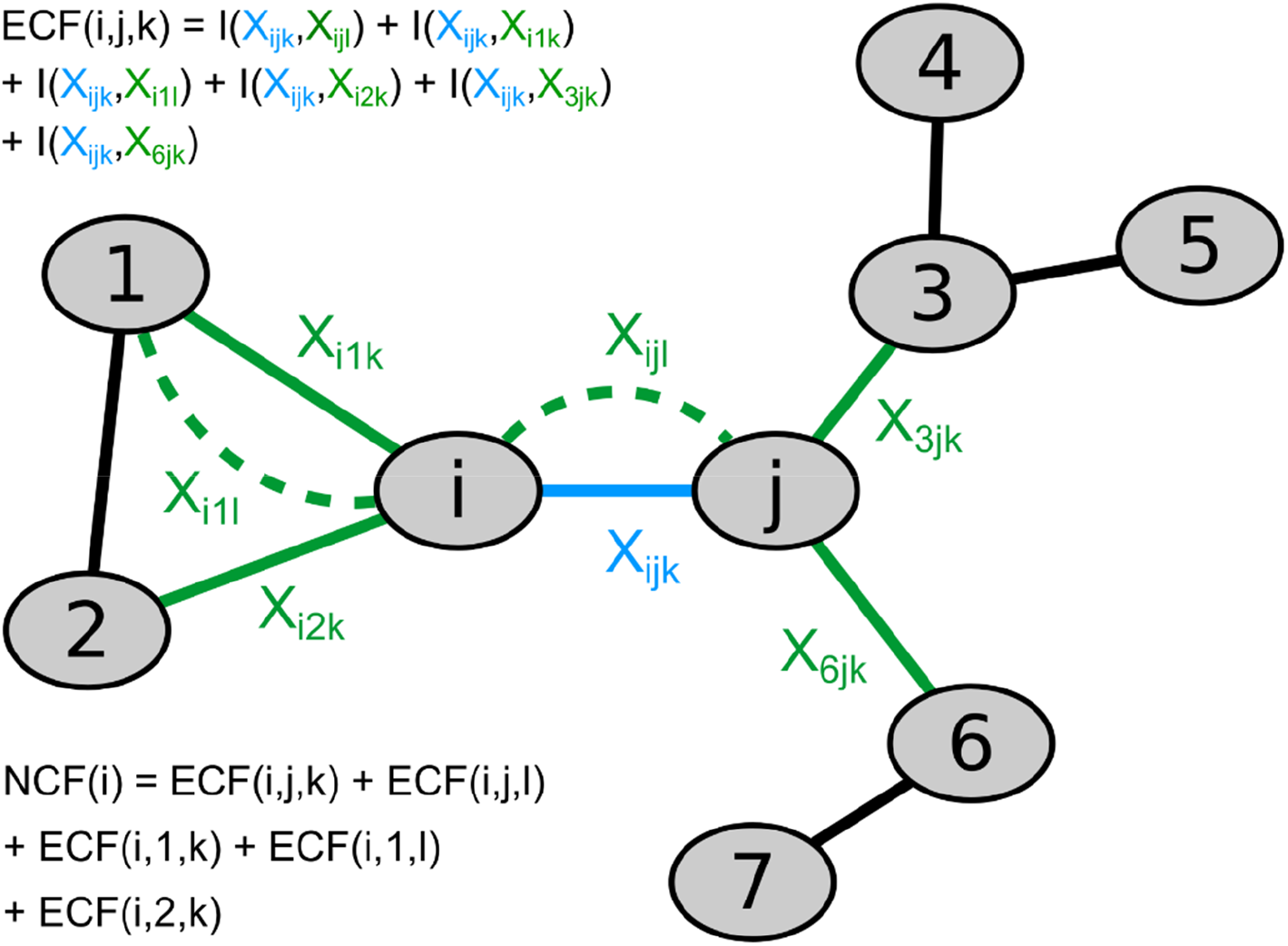
Example network demonstrating the calculation of edge correlation factor (ECF) and node correlation factor (NCF) scores. The ECF score of edge *i, j, k* (blue) is obtained by summing the mutual information of timeline *X*_*ijk*_ shared with the timelines of neighboring edges (green). The self-information *I*(*X*_*ijk*_, *X*_*ijk*_) is excluded. Subsequently, the NCF score of node *i* is calculated as the sum of ECF scores of all edges connected to *i*.

An essential feature of our model emerges from the definitions of the ECF, NCF and DNCF scores, namely the explicit locality of network effects. By limiting our analysis on the shared information between adjacent residues in the network, the influence of spurious correlation is reduced. To illustrate, consider that any pair of residues in a protein, no matter how far apart, would be compared. This would lead to a drastic increase of evaluated correlation terms, and thus more residue pairs showing high correlation by pure chance. At the same time, the probability that two residues influence each other directly in a substantial manner (i.e. without detectable changes in the residues between them) is lower if they are far apart, especially as the physical interactions included in our analysis, i.e. hydrogen bonds and carbon contacts, are of limited range. Adding up contributions of distant residues would thus substantially increase the noise introduced in the analysis. Instead, we propose that in most cases it is more productive to focus on the identification of neighboring residues directly exchanging information, and to analyze how they build chains of signaling residues. However, in instances of allosteric communication lacking this locality of effects, other methods may be more accurate.

### Molecular dynamics simulations

MD simulations in this work are based on the crystal structures of hPTP1E-PDZ2 in the apo state (PDB-ID: 3LNX) and bound to the C-terminal peptide of RA-GEF-2 (PDB-ID: 3LNY) as well as the corresponding solution NMR structures 3PDZ and 1D5G, using the first model provided in the files. These NMR structures were chosen to allow for direct comparison with previous studies (10, 39). Protein and ligand residues missing in the crystal structures were added based on their NMR structure analogues using Modeller 9.18 (52), creating 100 candidate structures and selecting the model with the best DOPE score for simulations and network analyses. MD simulations were performed using the Amber16-AmberTools17 software suite (53) with the Amber14SB force field (54) and TIP3P water (55). The system was solvated in a cubic water box using a minimum solute-face distance of 12 Å and 150 mM NaCl. For the nonbonded interactions a 12 Å direct space cutoff and PME summation for electrostatic interactions were applied. Energy minimization was performed until convergence to 0.01 kcal * mol^-1^ * Å^-1^ was reached using the XMIN minimizer. Afterwards, the volume of the solvent box was adjusted to a solvent density of 1.00 kg * m^3^. For all simulations a time step of 1 fs was applied and SHAKE (56) was used for hydrogen-containing bonds. Systems were gradually heated from 0 to 300 K over 1.7 ns using a variant of our published heatup protocol (57), restraining all heavy atoms by 2.39 kcal * mol^-1^ * Å^-2^ until 20 K and all backbone atoms until 200 K. For the first 1.2 ns of the heatup a Langevin thermostat was used with a collision frequency of 4 ps^-1^ and for the last 0.5 ns a Berendsen barostat was employed with a relaxation time of 2 ps. Afterwards the NPT ensemble was used with a slow coupling Berendsen thermostat at 300 K (coupling time: 10 ps) in combination with a Berendsen barostat (relaxation time: 5 ps). For each system, ten independent simulations were performed for 1 µs each (based on separate heatup runs and different randomized Langevin seeds). The initial 100 ns of each replicon were removed before analysis to reduce bias towards initial structures. Trajectory post-processing was performed with CPPTRAJ (58), using the “nativecontacts” command for contact timelines (saving both native and nonnative time series), and the “hbond” command for hydrogen bonds (distance cutoff 3.5 Å; angle cutoff 135°). Corresponding contact and hydrogen bond data from PDB structures were extracted with AIFgen.

### Protein structure networks

For analyses of protein structure networks and related quantities we used the SenseNet plugin (version 1.0.0) for Cytoscape (version 3.6.1) (32). CPPTRAJ outputs of contact and hydrogen bond analyses were extracted using AIFgen, a complementary tool encapsulating parts of SenseNet to generate networks on the command line. ECF scores were calculated with SenseNet using the therein implemented “Correlation” function set to the “Neighbor information” mode and summed to NCF scores using the “Degree” function. DNCF scores were calculated using the “Correlation” function set to “Neighbor information difference” followed by summation using the “Degree” function. As references for DNCF calculations (as in eq. 8), we selected the network generated from the corresponding ligand-bound simulation for the analysis of the network of the free protein, and vice versa. Contact betweenness centralities (BC) (59) and characteristic path length centralities (CPLC) (9) were calculated using the “Centrality” function of SenseNet (see S1 Text for details on these algorithms) and normalized using the min-max procedure. For high throughput analyses, we used the CyREST interface of Cytoscape to call the corresponding SenseNet functions. Plots were generated using matplotlib (version 3.0.3) (60) with pictures of molecular structures by VMD (1.9.3) (61) and open-source PyMOL (version 1.8.4.0) (62).

### Prediction of allosteric residues

Predictions were verified against methyl sidechain dynamics data (39), using classifications as allosterically active and inactive as defined by Cilia et al. (“NMR dataset”, n = 25, see S1 Table) (10). In that study, backbones of NMR structures and Monte Carlo sampling were used to find correlated side chain torsions. As this method was not applicable to alanine residues, the authors evaluated prediction performance using either the complete NMR dataset or a variant excluding alanine residues (“NMR-Ala dataset”, n = 21). To be consistent with these former studies, we chose to adopt this scheme in this work. Receiver Operating Characteristic (ROC) curves were generated by plotting, for various prediction score thresholds, the corresponding False Positive Rates (FPR) and True Positive Rates (TPR) with False Positives (FP), True Positives (TP), False Negatives (FN) and True Negatives (TN) according to the NMR datasets. In addition, we generated Precision-Recall (PR) curves based on Precision (PPV) and Recall (equivalent to TPR) scores. The overall prediction performance was evaluated by calculating the area under the curve for both ROC (rocAUC) and PR plots (prAUC) using trapezoidal integration.

## Results

### Features and Implementation of SenseNet

SenseNet reads interaction data from structure ensemble files in PDB format or MD trajectory analysis outputs generated by CPPTRAJ (58). By default, each node corresponds to a single amino acid and edges represent interactions on the amino acid level. SenseNet automatically determines the network topology from these timelines (Fig 2a), offering different adjustment options from removing rare interactions to considering only certain interaction types. Different levels of timeline analyses are possible, as users can either scroll through single time frames to investigate e.g. network evolution or time-dependent interactions, or analyze time-averaged networks. At any point during a running session, residue level nodes and associated interactions can be split into individual atoms, allowing for system specific tailoring of different resolution levels. As an example application providing a detailed demonstration of this concept, we refer to our previous study analyzing the recognition of different DNA modifications by the protein UHRF1 (63). SenseNet’s user interface is separated into the main network and three control areas (Fig 2b). The left panel allows access to implemented analysis functions and displays visualization status information, such as the selected edge weighting scheme or a bar to scroll through different time frames of the network. Whenever an analysis is performed, a summary of obtained results appears on the right panel, either as tables or plots. In addition, results are written into the node and edge data tables in the bottom region, from where they can be utilized by other analysis functions, either by SenseNet or other tools. This workflow, in combination with side-by-side network and structure visualization, allows for a rapid explorative cycle of performing quantitative analyses and intuitive exploration of the underlying structural details.

**Fig 2.**
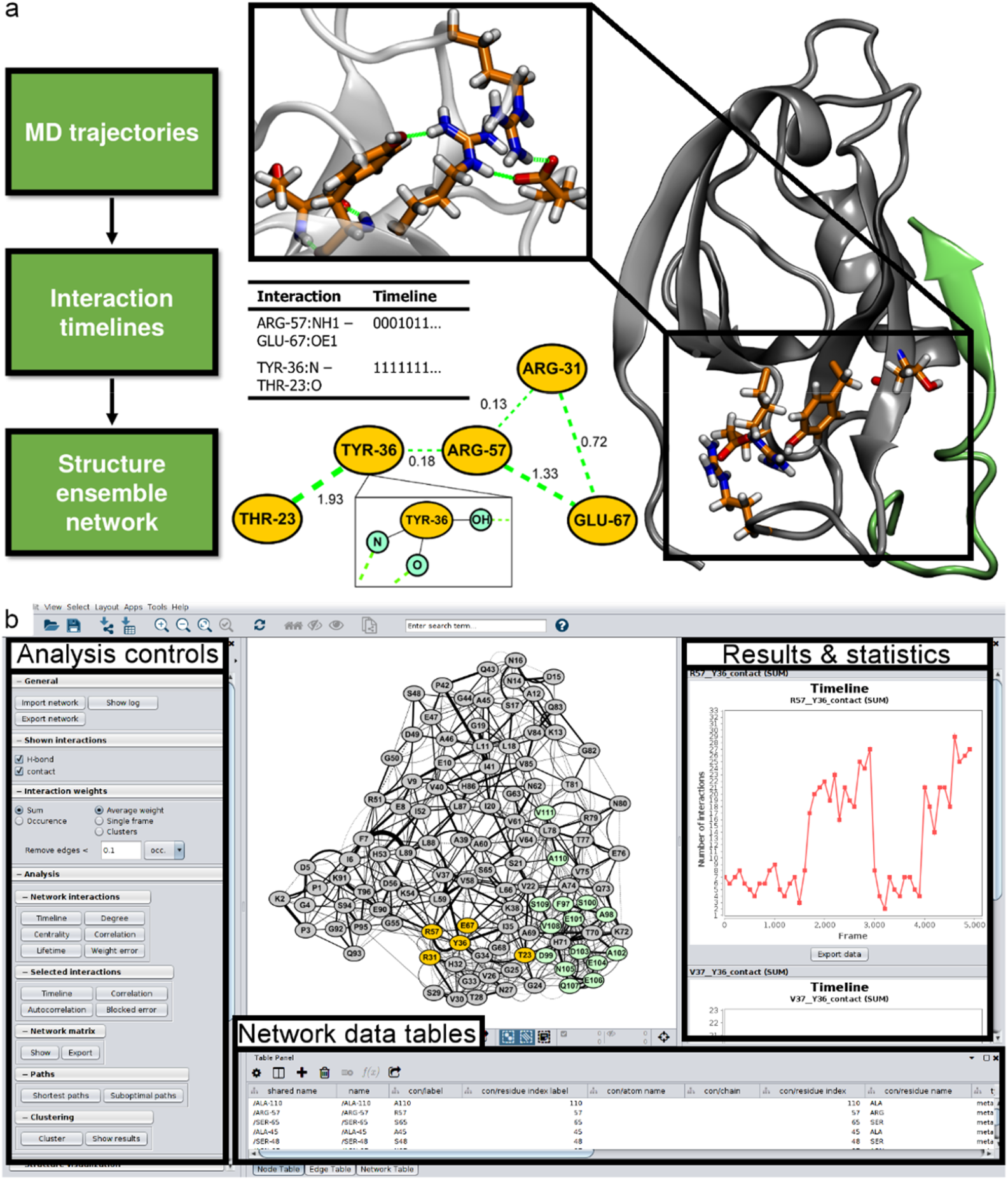
Example of parallel network and structure visualization using SenseNet. (a) Data representation, workflow and parallel representation of networks and molecular structures. (b) Example session showing the SenseNet GUI in Cytoscape.

For quantitative analysis of timeline data, SenseNet offers functions for calculating timeline correlation, entropy, autocorrelation, lifetime, clustering, and network comparison. In addition, search algorithms for shortest paths as well as centrality measures are provided. Analysis results are presented as tables or plots and can be exported as raw data or images. For large scale workflows, analyses can be automated via batch script files or the CyREST interface. Network and structure visualization can be carried out in parallel by connecting SenseNet to the PyMOL (62), VMD (61), or UCSF Chimera (64) structure viewers, automatically highlighting selected nodes and edges from the network in the protein structure.

### Evaluation of different methods for allosteric prediction

First, we reinvestigated the allosteric prediction performance of betweenness centralities (BC) and characteristic path length centralities (CPLC) based on networks generated from NMR and crystal structures, which had previously shown poor prediction performance for the PDZ2 system with CPLC as the best performing centrality model (10). This allowed us to verify our implementation and to compare different network methods based on the same dataset. In line with the aforementioned work, we determined ROC and PR curves measuring the prediction accuracy of tested models with respect to the NMR dataset, which is composed of allosteric and non-allosteric residues based on methyl sidechain dynamics, and the corresponding NMR-Ala dataset variant excluding alanines (10, 39) (S1 Table). In an attempt to replicate the network centrality predictions from Cilia et al. (NMR: 0.54, NMR-Ala: 0.59) (10), we calculated CPLC scores based on the crystal and NMR structures of the PDZ2-RA-GEF-2 complex using a carbon contact distance cutoff of 5 Å. For the NMR structure, resulting rocAUC scores were very close to the previously reported values (NMR: 0.55, NMR-Ala: 0.56) and only modestly higher for the crystal structure (NMR: 0.65, NMR-Ala: 0.69), indicating that the differences are only due to subtly differing details in network implementations.

In contrast to the centrality approach, interaction timelines generated from structure ensembles allow to additionally analyze the correlation between interactions, as quantified by the NCF and DNCF scores (see Materials & Methods). In general, residues with high NCF scores provide information, through linear and nonlinear correlation, about the interaction state of their environment. While the NCF estimates the information of residues within a single simulation, the DNCF score models the corresponding differences between two simulations, e.g. with and without a ligand. In order to obtain the structure ensembles necessary for calculation of these scores, we performed ten 1 µs MD simulations of the free PDZ2 domain and the PDZ2-RA-GEF-2 peptide complex. Timelines of contacts and hydrogen bonds were extracted and converted into protein structure networks using AIFgen and analyzed using SenseNet. First, we systematically evaluated all compared network methods (BC, CPLC, NCF, DNCF) using a grid search of 48 parameter combinations (S2 Table). These combinations were obtained by varying the contact distance cutoff from 4 to 9 Å, the interaction subset settings (all or only inter-sidechain interactions), and networks generated from different sources (apo- or peptide-bound structures; NMR or crystal structures). To understand which parameters are most important for prediction performance, we grouped all data points according to these categories followed by analysis of the obtained rocAUC score distributions. In the following, we focus predominantly on the results obtained for the NMR-Ala dataset, as alanine residues proved to be particularly difficult to predict for all methods tested here as well as those previously published. Fig 3a shows that average rocAUC scores over all combinations were consistently highest for the DNCF method, followed by NCF and finally CPLC and BC, which registered 8 – 11 % lower average AUC scores compared to the former methods. In a more detailed view (Fig 3b), we observed that on average, prediction performances improved if apo PDZ2 was used as starting structure compared to peptide bound systems, with relatively small differences for CPLC, BC, and DNCF (up to 5 %), but more substantial improvements for NCF (up to 9 %). Interestingly, the NCF prediction performance based on the apo systems was almost as high as the DNCF scores although, in contrast to DNCF, they do not contain any information about the ligand. Regarding the set of included interactions in the network (Fig 3c), rocAUC scores increased on average by 2 - 4 % if only inter-sidechain interactions were considered. Finally, analysis of contact cutoff distances shows that BC and CPLC method performances appear to peak at 6 Å, whereas a 4 to 5 Å cutoff worked best for the DNCF and NCF methods (Fig 3d). Many of the observed trends are reflected, to a lesser degree, on the full NMR reference set which includes alanine residues (S1 Fig). In conclusion, we first observe that all parameter categories follow consistent trends, highlighting the importance of parameter choice for prediction quality. Second, this consistency is also observed if the different methods are compared, i.e. the favorable performances of NCF and DNCF models relative to centralities are reflected throughout all parameter settings.

**Fig 3.**
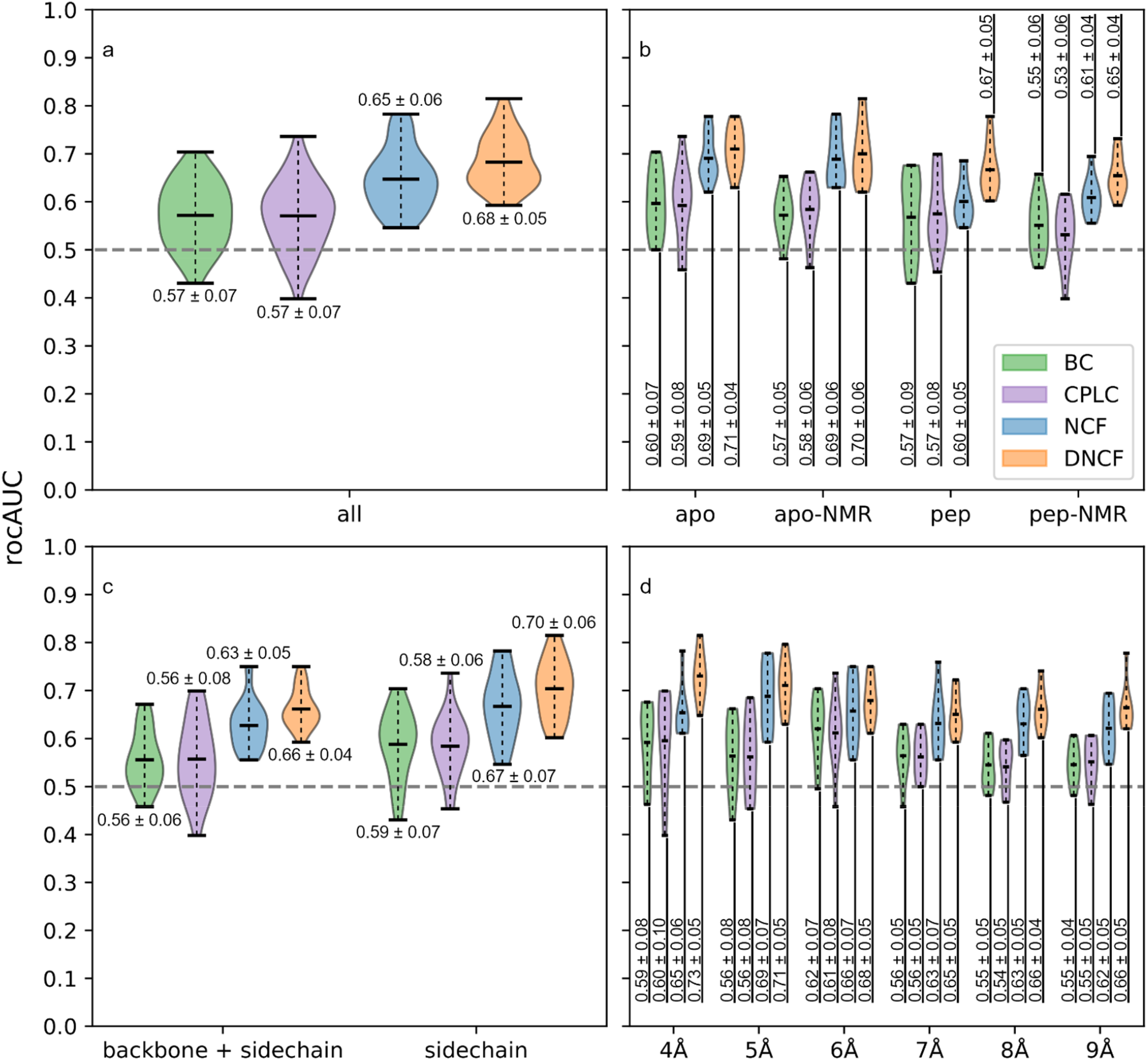
Influence of network parameters on prediction model performance based on the NMR-Ala reference set. Shaded areas show distribution estimates based on a gaussian kernel with added labels for mean and standard deviation. (a) Distributions including all parameter combinations. (b) Source of analyzed network data: Crystal structures (apo, pep) or NMR based structures (apo-NMR, pep-NMR). (c) Interaction subset: All interactions or sidechain-exclusive networks. (d) Distance cutoff for carbon-carbon contacts in the network.

The best performing CPLC model was obtained for the apo PDZ2 crystal structure and a carbon contact cutoff of 6 Å in a sidechain exclusive network, interestingly differing from the original evaluation discussed above (5 Å and including backbone interactions) (10). Using the optimized parameters, the rocAUC score for the NMR-Ala dataset increased by 5 % to 0.74, while performance for the NMR dataset degraded by 1 % to 0.64, respectively (Table 1). The corresponding prAUC scores increased by 2 % for the NMR dataset (0.75 to 0.77) and 5 % for NMR-Ala (0.78 to 0.83). The BC method performed optimally with the same parameter set as CPLC, but with about 3 to 4 % lower rocAUC scores (Table 1). Overall, only modest performance improvements could be achieved for the CBC and CPLC methods by variation of network parameters.

**Table 1.**
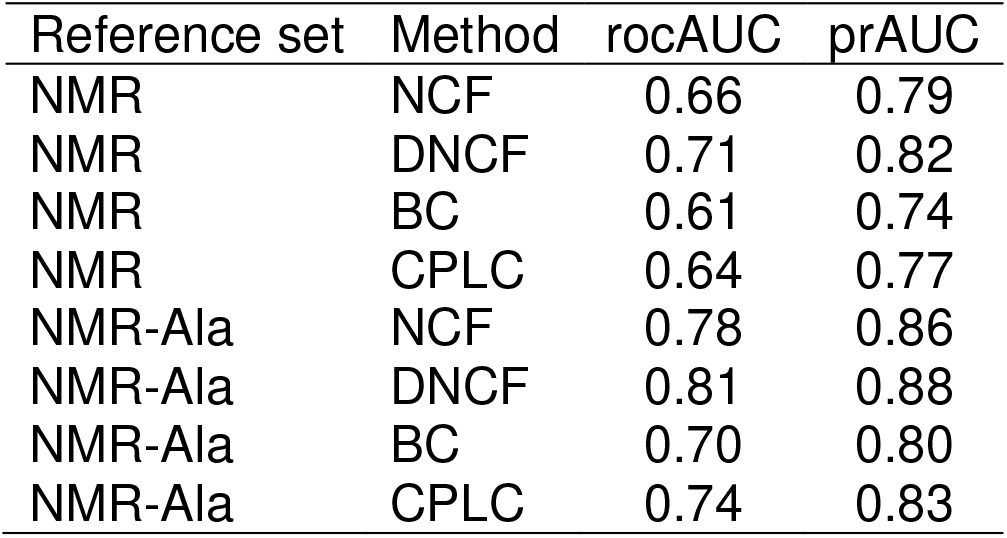
Allosteric prediction performance of network-based models.

For both DNCF and NCF models, the optimal parameter set consisted of a 4 Å contact cutoff in a sidechain exclusive network using simulations of the apo-NMR PDZ2 structure. Of all settings tested in the parameter search, DNCF was found to be the best overall predictor, achieving a rocAUC of 0.71 and prAUC of 0.82 on the full NMR set, which corresponds to a 5 to 7% improvement compared to the CPLC model. Accordingly, the performance on the NMR-Ala set was also higher than for the centrality methods with a rocAUC of 0.81 and a prAUC of 0.88. The best NCF model showed similar overall trends, but individual AUC scores were 1 – 5 % lower (Table 1). In line with most published methods, rocAUC scores were consistently 7 – 10 % lower for the NMR dataset compared to NMR-Ala, which highlights the general difficulty for predicting this residue type (Table 2).

**Table 2.** Comparison of DNCF prediction performance with other published computational methods.

| Reference set | Method | rocAUC | prAUC |
| --- | --- | --- | --- |
| NMR | DNCF | 0.71 | 0.82 |
| NMR | NMR/MC | 0.74 | 0.82 |
| NMR | RRS | 0.65 | 0.75 |
| NMR | REDAN | 0.67 | 0.65 |
| NMR-Ala | DNCF | 0.81 | 0.88 |
| NMR-Ala | NMR/MC | 0.81 | 0.87 |
| NMR-Ala | RRS | 0.72 | 0.80 |
| NMR-Ala | REDAN | 0.62 | 0.61 |

It has been pointed out that the allosteric residue sets from published computational predictions differ substantially for the PDZ2 system (41), fueling our interest determining how well these models agree with the NMR datasets. However, comparing models based on binary classifications alone can be misleading, since each classification relies on an implicit sensitivity threshold which might differ drastically between models. ROC and PR curves are more suitable for this task since they evaluate prediction performances at all possible thresholds, but require raw prediction scores, which are not always available. Fig 4 shows the ROC and PR curves for the models described above and those for which accompanying literature included the necessary scores. We observed comparably high performances for the DNCF and NMR/MC (10) models (Table 2, differences within 1 – 2 %), followed by RRS (50) and REDAN (46). As the NMR/MC model requires NMR structure data, the DNCF method offers a substantial advantage as the necessary simulations can be based on much more commonly available crystal structures. Thus, although these two methods show comparable accuracy, we expect that the DNCF method can applied to a wider range of systems. We also believe that the method has the potential to show improved results for systems for which induced fit phenomena are important, i.e. for which the conformational ensembles of the apo- and holo-structures differ considerably.

**Fig 4.**
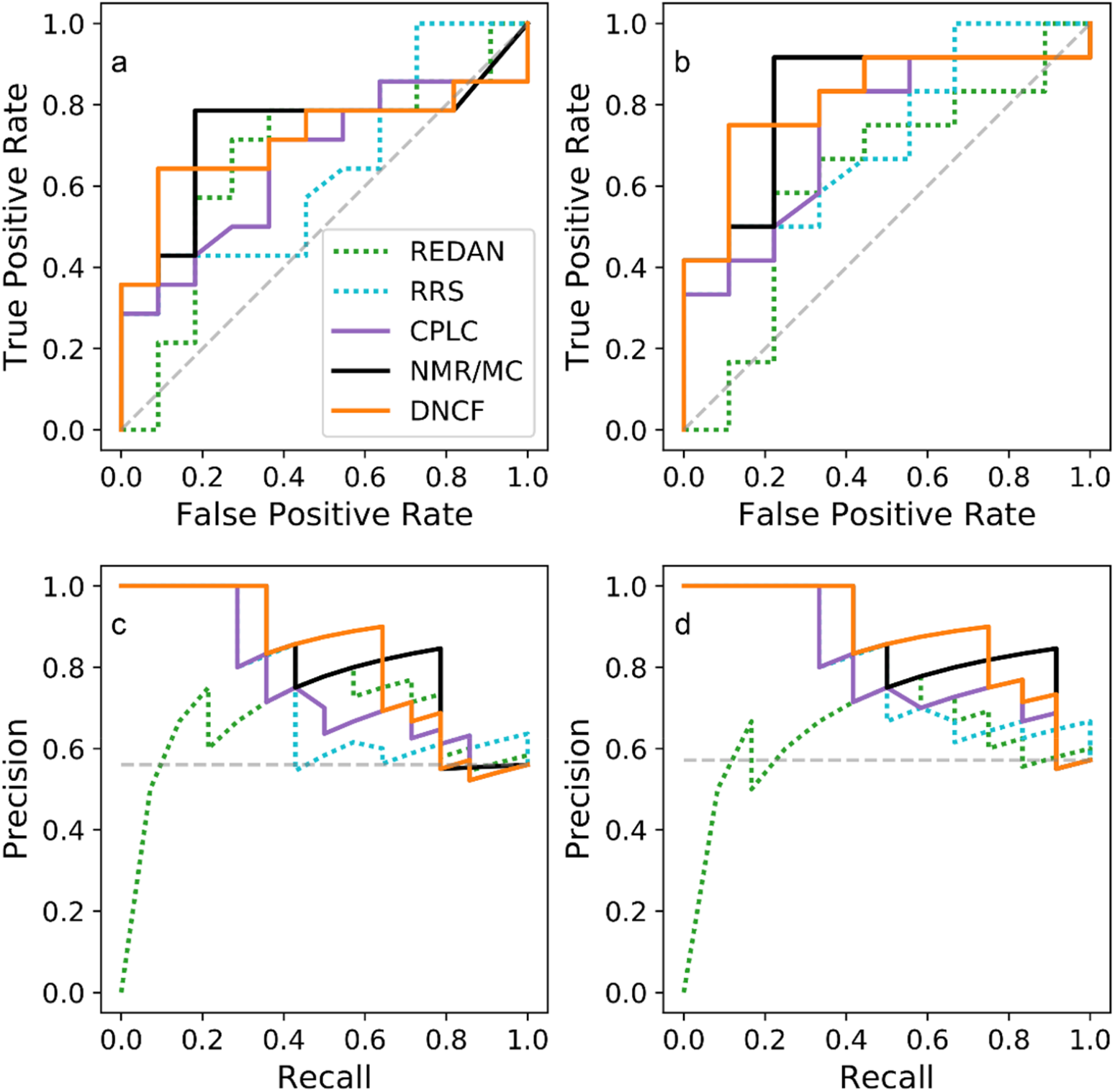
ROC and PR curves of selected prediction models. (a) ROC curve based on the NMR reference set. (b) ROC curve based on the NMR-Ala reference set. (c) PR curve based on the NMR reference set. (d) PR curve based on the NMR-Ala reference set.

### Prediction of allosteric residues in the hPDZ2 domain

Having established good agreement between DNCF scores and allosteric residues, we investigated the usefulness of these additional features for the biochemical interpretation of our predictions in the PDZ2 structure. Integrating the DNCF scores of the model described above into the structure network (Fig 5a,b) reveals two high scoring clusters of residues (clusters I and II). The majority of allosteric residues of the NMR dataset are located in cluster I, which stretches from the top region of the binding pocket towards helix α1 and sheet β1 (Fig 5b-d). On the other hand, cluster II encompasses the lower part of the binding pocket surrounding the flexible loop L1 (residues 24-33), including the allosteric residues V26 and V30, furthermore its interaction partners R57, Y36, and finally the C-terminal region.

**Fig 5.**
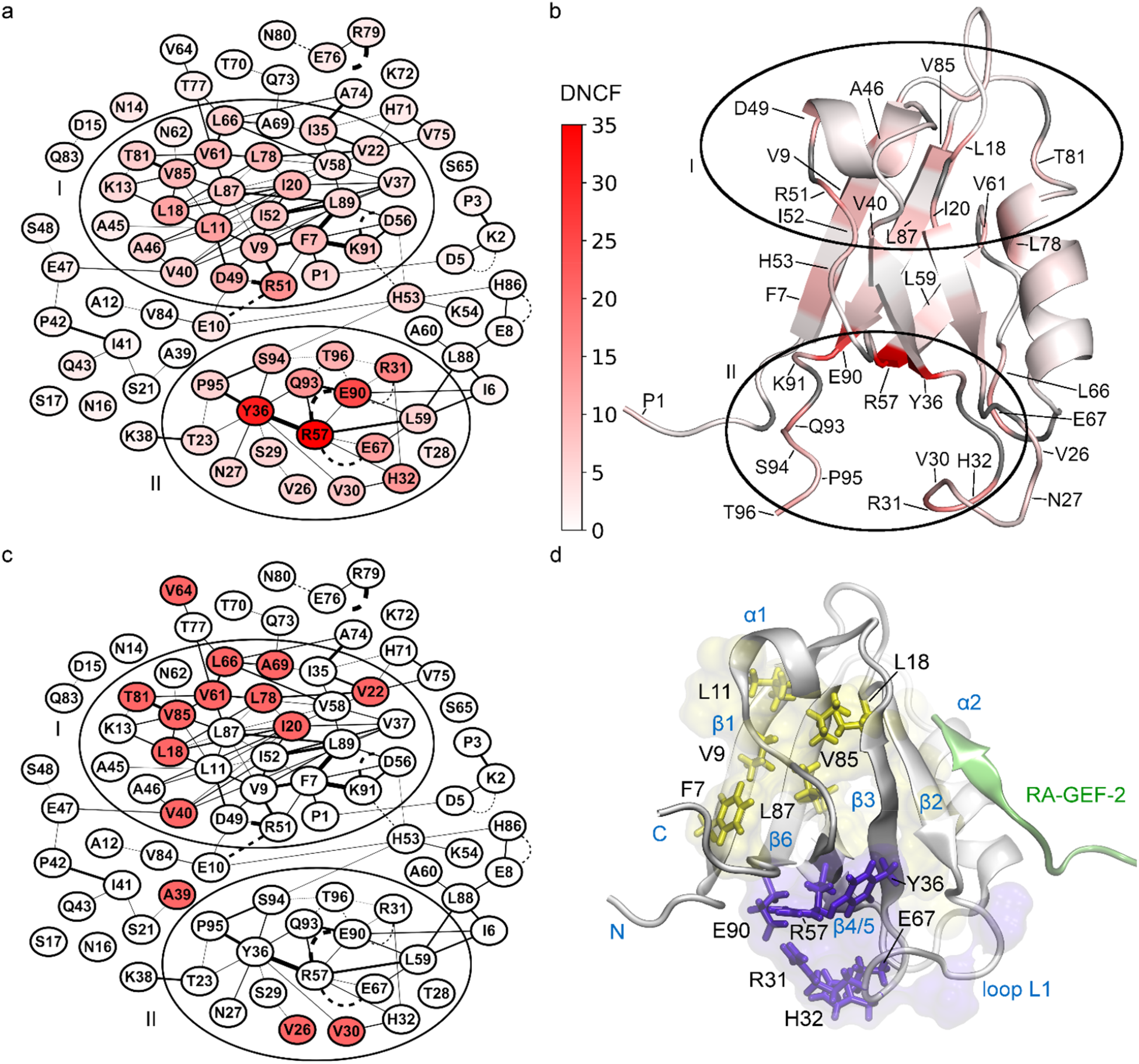
Allosteric predictions of the final DNCF model mapped to PDZ2 structures. For visual clarity, only edges occurring in ≥ 0.1 % of simulation time are shown. (a) Network representation of DNCF predictions. Nodes are colored from low (white) to high (red) DNCF scores. (b) DNCF scores mapped to the apo PDZ2 structure (PDB-ID: 3PDZ). (c) Network showing experimentally determined allosteric residues (red) from the NMR dataset. (d) Allosteric clusters mapped to the RA-GEF-2 bound PDZ2 structure (PDB-ID: 1D5G): Cluster I (yellow surface) and Cluster II (purple surface). Specific residues discussed in the text are additionally shown as sticks.

Comparing these observations to other network scoring methods, the NCF model shows a very similar cluster structure (Fig 6a), whereas for CPLC we observed increased scores for residues located next to the peptide binding groove, e.g. V22, L66, H71, A74, V75 and L78 (Fig 6b-d). This can be explained directly by the definition of CPLC (S1 Text), which attributes high scores to residues bridging structural modules, e.g. binding grooves. On the other hand, centrality scores for loop L1 in cluster II are substantially lower than in the timeline-based NCF and DNCF methods, which might be explained by the difficulties of a single structure network to represent the switching contacts of flexible regions. This indicates that centrality methods may fail to account for regions with intrinsic flexibility like the L1 loop, for which methods based on structure ensembles are potentially more appropriate.

**Fig 6.**
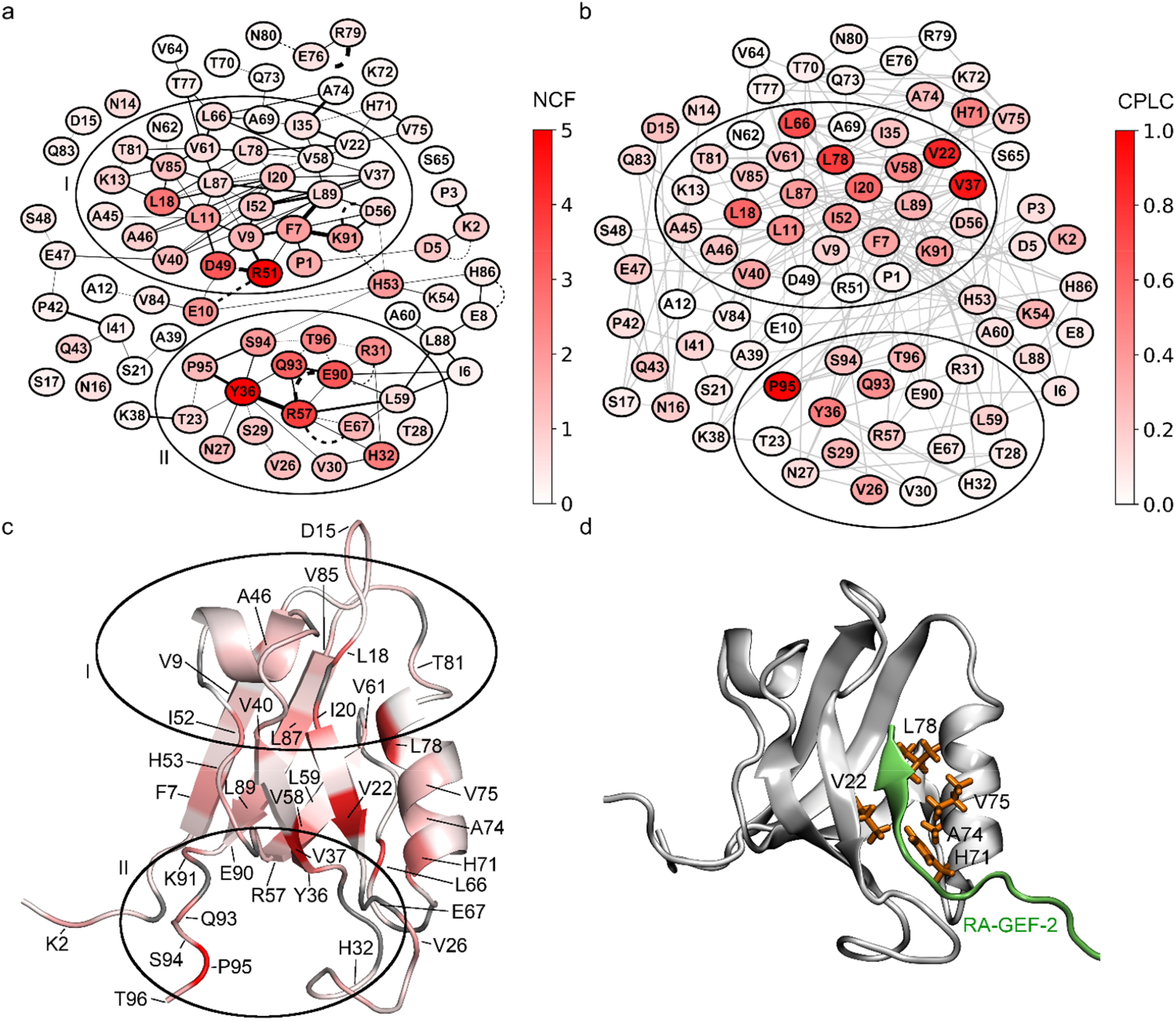
Allosteric predictions of the final NCF and CPLC models mapped to PDZ2 structures. Nodes colored from low (white) to high (red) scores. (a) Network representation of NCF predictions. For visual clarity, only edges occurring in ≥ 0.1 % of simulation time are shown. (b) Network representation of CPLC predictions. Edge colors are shown in light grey to increase clarity. (c) CPLC scores mapped to the apo PDZ2 structure (PDB-ID: 3PDZ). (d) Notable residues predicted by CPLC mapped to the RA-GEF-2 bound PDZ2 structure (PDB-ID: 1D5G).

### Consensus model of allosteric information flow in PDZ2

Finally, we defined a new consensus model of allosteric information flow consolidating our and previous prediction models. For this we first determined a “consensus set” composed of residues predicted as allosteric in ≥ 50 % from a selection of published studies (S3 Table) (10, 42-45, 47-51, 65). Next, we obtained a core set of allosteric candidates from our DNCF model, using the score threshold closest to the top left corner in Fig 4b (6.17 bits in S4 Table; TPR: 0.75; FPR: 0.11). This core prediction set (Fig 7 and S5 Table) contains 9 out of 14 residues from the NMR dataset and 11 of the 18 from the consensus set, while 14 residues are complementary predictions. Of these infrequently predicted residues, three form a contiguous surface located on the sheet β1 (F7, V9, L11), connected via L18, V85, and L87 to the peptide binding pocket (Fig 5d). In NMR experiments, V9 was shown to respond to the binding pocket I20F mutation with L11 and L87 as presumed linker residues (40), an interpretation supported by our model. Notably, the clusters surrounding V9 and Y36 agree very well with the DS3 and DS4 regions described previously (10). Predictions of the C-terminal tail residues (93 to 96) are difficult to assess as the high flexibility of free chain termini might not properly represent the common biological state, i.e. PDZ2 embedded in a multi-domain protein. Previous studies have formulated the idea of up to four separate distal sites (DS1 - DS4) identified by following the interconnected surfaces of allosteric residues (10, 39, 65). Our results suggest the existence of at least two allosteric clusters: Cluster I which encompasses DS1, DS2, and DS3, while cluster II corresponds to DS4.

**Fig 7.**
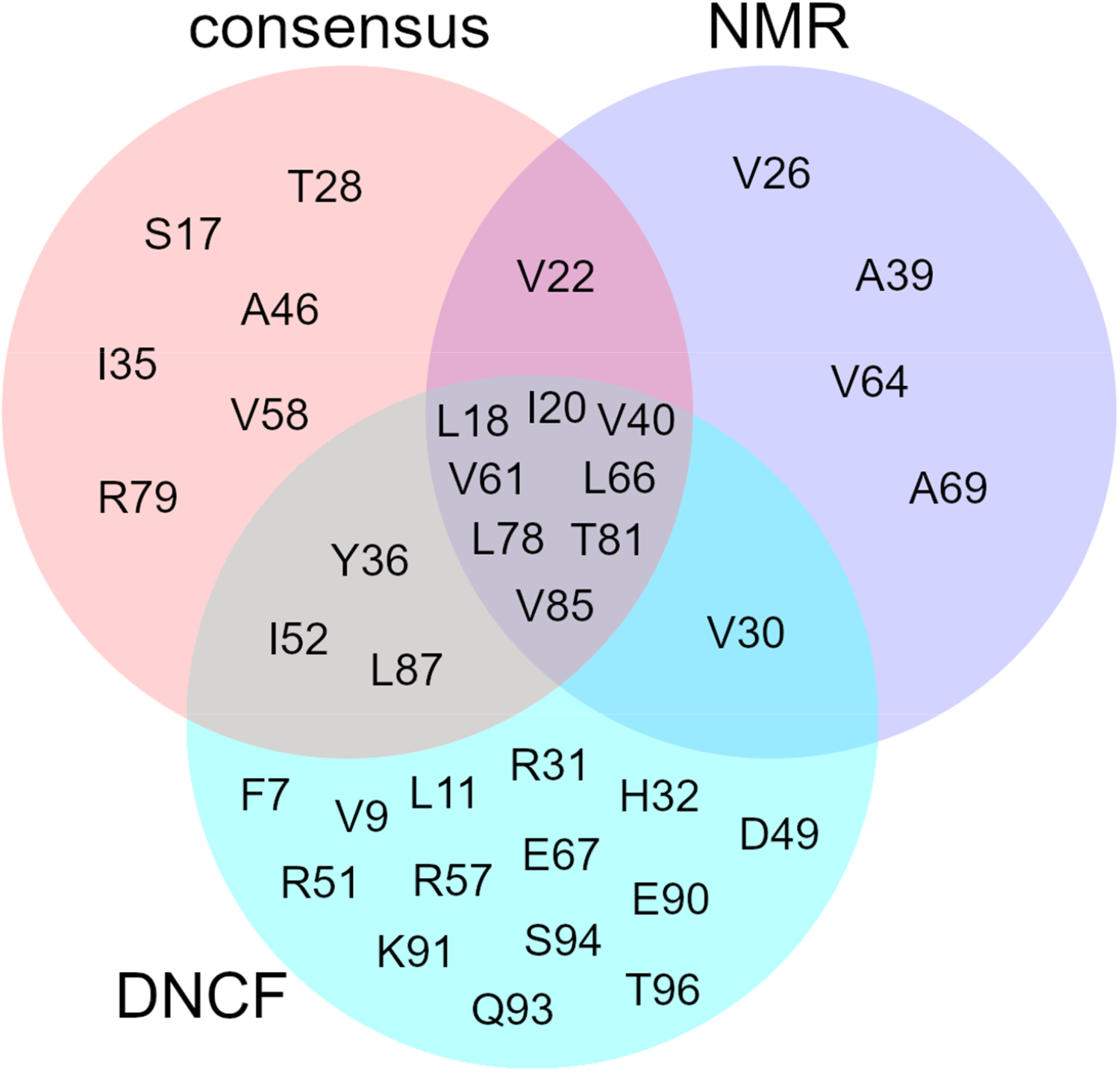
Intersection of the DNCF allosteric core set, NMR reference set, and the computational prediction consensus set.

## Discussion

Integration of interaction timelines from molecular dynamics simulations into protein structure networks provides a promising framework for investigating dynamic effects in proteins such as allostery. In this work, we introduce our network analysis tool SenseNet which builds on this theoretical foundation. Using the PDZ2 domain as a reference system, we evaluated four allosteric prediction models implemented in SenseNet, i.e. BC, CPLC, NCF and DNCF, and determined a set of network parameters optimizing their accuracy. Our results are consistent with literature data, as structure networks frequently use carbon contact cutoff distances between 4 - 6 Å (10, 19, 47, 66, 67), which corresponds approximately to the upper limit of attractive Van-der-Waals interactions. The trend for better prediction results using apo protein states might reflect the observed rigidification of the ligand binding site after binding (39) and is in line with previous suggestions that allosteric mechanisms may be intrinsic properties of apo structures (42, 68). Finally, the improvements observed in sidechain exclusive networks mirror the origins of the NMR dataset, which was obtained from methyl sidechain dynamics (39). This also highlights an important caveat for comparing prediction models, as some methods might by design match certain types of experimental data more closely than others. The final allosteric model, based on the DNCF method, was found to be one of the models aligning most closely to experimental data alongside NMR/MC. However, the DNCF approach offers three distinct advantages to NMR/MC: First, MD simulations for DNCF analyses can be started from only a single, e.g. X-ray, structure, while NMR/MC needs an NMR structure ensemble, which are far rarer and more limited to small proteins. Second, the DNCF method includes all residue types, while NMR/MC by definition cannot predict alanine residues. Third, the DNCF method has the potential to detect induced fit-based conformational changes, which are often not directly detectable in the structural ensembles of the apo-state alone. Mapping the results of our DNCF model to the structure of PDZ2 suggests the protein contains two distinct allosteric sites. Most of the experimentally verified allosteric residues from the NMR dataset are located in cluster I, while cluster II has little support from the experimental dataset as the region encompasses only four residues with methyl groups. To fill this gap, alternative experiments may be necessary such as mutational studies connected to changes in PDZ mediated activation. The locations of our observed clusters are matched by several other computational predictions (42, 43, 45). Nevertheless, our data contrasts with studies reporting up to four distinct allosteric sites (10, 39, 65) by suggesting that these four sites are partially overlapping, leaving only two clearly separated allosteric regions. The variance in published allosteric predictions in the PDZ2 domain may be explained by the fact that the experimentally verified data in a single protein are naturally sparse, leading to potentially large error margins for validation. In addition, for many cases quantitative scores are not reported along binary classifications, impeding direct comparison of predictions. To improve prediction models, large scale studies including multiple proteins, computational methods, and experimental data sources will be necessary. With SenseNet we provide a network analysis tool offering considerable advantages over existing implementations: First, by defining edges via interaction timelines, all conformational states of a simulation are readily available for analysis, which is not possible if interactions are reduced to correlation coefficients. Second, adopting a multi-resolution approach via mapping of sub-structures of varying sizes to nodes (from atoms to residues) allows the creation of application-specific network topologies that reduce the underlying structural differences to the most informative level of details. Finally, integration of our tool into Cytoscape allows users to complement their analyses with the community driven ecosystem of biological network analysis plugins, e.g. by connecting structural analysis with system biological or sequence/evolutionary information. Based on these concepts, SenseNet provides an analysis platform implementing a range of well tested analysis algorithms, an easy-to-use UI driven implementation, and interactive side-by-side structure visualization. Together, these features serve as a potential foundation for wide application of timeline-based protein structure networks, paving the way for comparative studies to improve model accuracies and aid experiments in unveiling detailed mechanisms of dynamic processes in biomolecules.

## Supporting Information captions

**S1 Text. Measures of centrality in protein structure networks**.

**S1 Fig. Influence of network parameters on prediction model performance based on the NMR reference set**. Shaded areas show distribution estimates based on a gaussian kernel with added labels for mean and standard deviation. (a) Distributions including all parameter combinations. (b) Source of analyzed network data: Crystal structures (apo, pep) or NMR based structures (apo-NMR, pep-NMR). (c) Interaction subset: All interactions or sidechain-exclusive networks. (d) Distance cutoff for carbon-carbon contacts in the network.

**S1 Table. NMR reference set of experimentally verified allosteric and non-allosteric residues**. Allosteric residues are represented by a value of 1, non-allosteric residues by a value of 0.

**S2 Table. Prediction model performances for all tested network parameter combinations**.

**S3 Table. Computational predictions of allosteric residues including the DNCF model and previously published methods**.

**S4 Table. Residue scores of final DNCF, NCF, and CPLC models**.

**S5 Table. Comparison of the DNCF allosteric core set with the NMR reference and computational prediction consensus sets**.

**S1 File. Initial structures, topologies, and input files for molecular dynamics simulations**.

