## Supporting Text and Figures for "SenseNet, a tool for analysis of protein structure networks obtained from molecular dynamics simulations"

### Supporting Information

#### S1 Text. Measures of centrality in protein structure networks

Various centrality models have been proposed to quantify the importance of nodes in the context of the overall network topology. In this work, we use two centrality variants commonly applied to protein structure networks. Betweenness centrality (BC) (1-3) measures the fraction of shortest paths a residue contributes to and is defined as

$$BC(i) = \sum_{j,k \in N,\ i \neq j \neq k} \frac{\sigma_{jk|i}}{\sigma_{jk}} \tag{1}$$

where $i, j, k$ belong to the set of nodes $N$, $\sigma_{jk}$ is the number of shortest paths between $j$ and $k$, and $\sigma_{jk|i}$ is the number of shortest paths between $j$ and $k$ passing through $i$. In contrast, characteristic path length centrality (CPLC) (4) is based on the effect of a node on the characteristic path length, i.e. the average length of shortest paths in the network

$$L = \frac{1}{N_p} \sum_{i,j \in N,\ i > j} d(i,j) \tag{2}$$

where $N$ is the set of nodes, $N_p$ is the number of node pairs in the network and $d(i,j)$ is the minimum number of edges to be traversed between $i$ and $j$. The CPLC score corresponds to the effect of removing a node on the characteristic path length of the network, which can be expressed as

$$CPLC(i) = |L - L_i| \tag{3}$$

where $L_i$ is the characteristic path length of the network after removal of node $i$.

#### Figures und Tables

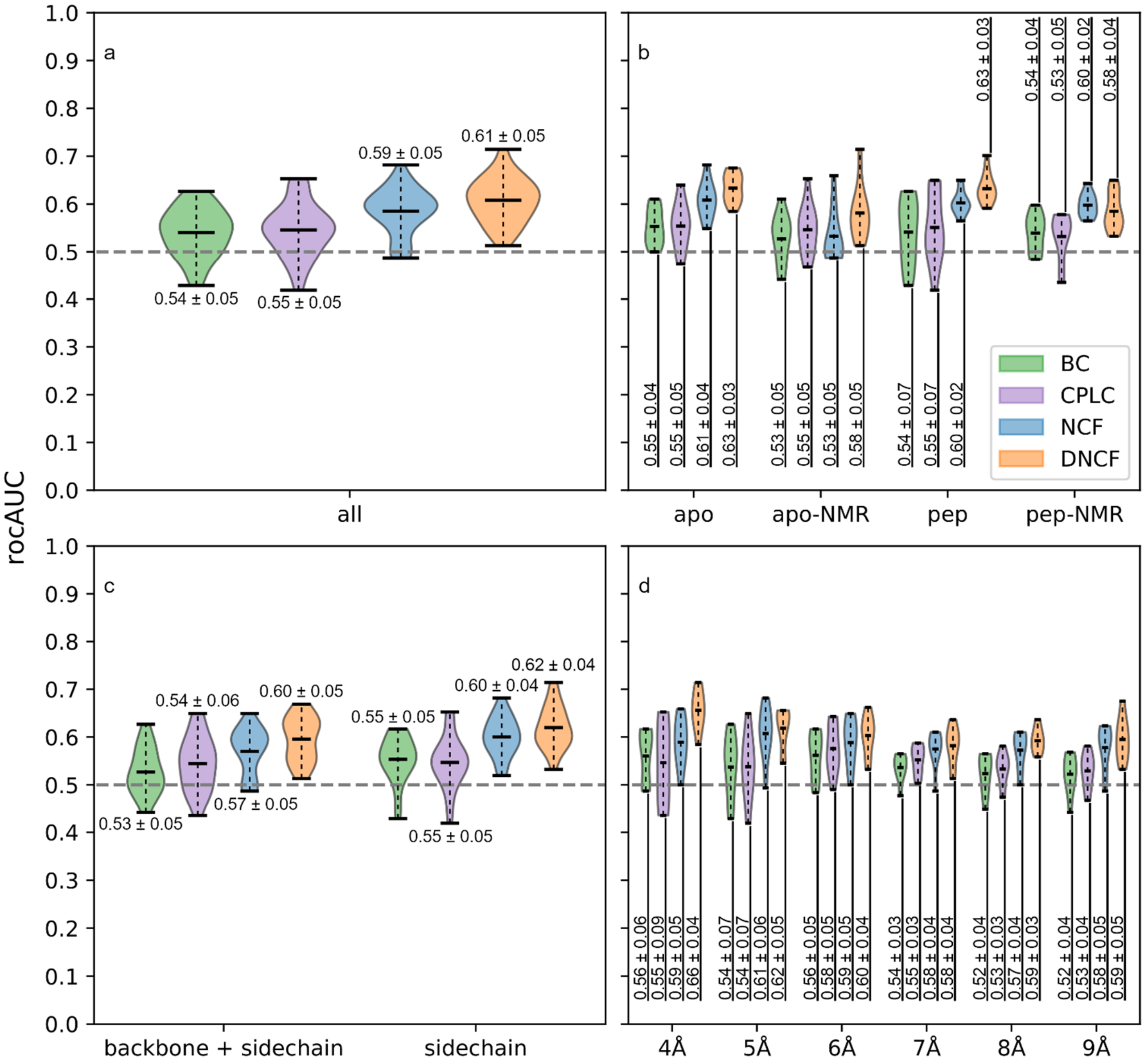


**S1 Fig. Influence of network parameters on prediction model performance based on the NMR reference set.** Shaded areas show distribution estimates based on a gaussian kernel with added labels for mean and standard deviation. (a) Distributions including all parameter combinations. (b) Source of analyzed network data: Crystal structures (apo, pep) or NMR based structures (apo-NMR, pep-NMR). (c) Interaction subset: All interactions or sidechain-exclusive networks. (d) Distance cutoff for carbon-carbon contacts in the network.

**S1 Table. NMR reference set of experimentally verified allosteric and non-allosteric residues**. Allosteric residues are represented by a value of 1, non-allosteric residues by a value of 0.

**S2 Table. Prediction model performances for all tested network parameter combinations.**

**S3 Table. Computational predictions of allosteric residues including the DNCF model and previously published methods.**

**S4 Table. Residue scores of final DNCF, NCF, and CPLC models.**

**S5 Table. Comparison of the DNCF allosteric core set with the NMR reference and computational prediction consensus sets.**

**S1 File. Initial structures, topologies and input files for molecular dynamics simulations.**
