## Supporting File 1 for "SenseNet, a tool for analysis of protein structure networks obtained from molecular dynamics simulations": README

Input files for molecular dynamics simulations for use with AMBER

Software used: 
Amber16/AmberTools17 (https://ambermd.org/)

Files and directories included in the manuscript Supplemental Information .zip archive:
- em: AMBER Input files for energy minimization
- heatup: AMBER Input files for heatup runs
- prod: AMBER Input files for production runs
- structures: Input structure and topology files
-- .incrd Initial structure for simulation (AMBER format)
-- .pdb Initial structure for simulation (PDB format)
-- .prmtop Topology file for simulation (AMBER format)

Files deposited in the Dryad data repository (doi:10.5061/dryad.3r2280gf2 - not active during peer review; please use https://datadryad.org/stash/share/CMlMjFaL1xw_ThaDl2DjqiwPuT7oJuJx3jU5VBc9XNk)
- .prmtop Topology file for analysis
- .nc Trajectory coordinate files used for analysis (AMBER Netcdf format). All replicas were merged (concatenated) into a single trajectory file.
